# Single-molecule FISH in Drosophila muscle reveals location dependent mRNA composition of megaRNPs

**DOI:** 10.1101/156091

**Authors:** Akiko Noma, Carlas S. Smith, Maximillian Huisman, Robert M. Martin, Melissa J. Moore, David Grunwald

## Abstract

Single-molecule fluorescence in-situ hybridization (smFISH) provides direct access to the spatial relationship between nucleic acids and specific subcellular locations. The ability to precisely localize a messenger RNA can reveal key information about its regulation. Although smFISH is well established in cell culture or thin sections, methods for its accurate application to tissues are lacking. The utility of smFISH in thick tissue sections must overcome several challenges, including probe penetration of fixed tissue, accessibility of target mRNAs for probe hybridization, high fluorescent background, spherical aberration along the optical axis, and image segmentation of organelles. Here we describe how we overcame these obstacles to study mRNA localization in *Drosophila* larval muscle samples that approach 50 μm thickness. We use sample-specific optimization of smFISH, particle identification based on maximum likelihood testing, and 3-dimensional multiple-organelle segmentation. The latter allows using independent thresholds for different regions of interest within an image stack. Our approach therefore facilitates accurate measurement of mRNA location in thick tissues.

## Introduction

Localization of mRNAs is a central mechanism for regulating mRNA function (Wu et al., 2016). Single-molecule fluorescence in situ hybridization (smFISH) provides snapshots of individual mRNA distribution patterns throughout cells. Initial smFISH protocols used long (50-nt) hybridization probes uniformly labeled with ~5 dyes (Femino et al., 1998). Because it introduces fluctuations in signal strength, however, non-uniform labeling of long probes decreases the threshold of detection, especially for RNA clusters. More recently, the smFISH protocol was simplified to use shorter (20-nt) probes labeled with a single fluorophore (Raj et al., 2008). These probes are cheaper to make and easier to label. Moreover, multiplexing up to 48 probes against a single mRNA species yields high contrast between specific and nonspecific binding (Raj et al., 2008).

Typically, smFISH is used to detect individual mRNAs in cultured cells — e.g., yeast cells, monolayer cultures, or isolated neurons (Buxbaum et al., 2014; Coassin et al., 2014; Kwon, 2013; Shaffer et al., 2013). Yet many cell types, such as multi-nucleate muscle cells, lose key tissue-specific features in single cell cultures. Within muscle tissue, the relevant dimensions of cells and nuclei make it desirable to work with thick (>40 μm) tissue sections. However, available smFISH protocols for tissues, including *Drosophila* imaginal wing discs and *C. elegans* larvae and embryos (Lyubimova et al., 2013; Oka and Sato, 2015; Raj et al., 2008) are limited to 5 to 10 μm sections. Performing smFISH in larger tissue volumes poses seven specific challenges: (1) fixatives and probes must penetrate deep into the tissue and into both cytoplasmic and nuclear compartments, but fixative and probe penetration rates differ between the cytoplasm and nucleus and from sample to sample; (2) probes must be able to efficiently access target mRNAs, but mRNAs are structured and coated with proteins so hybridization sites can be masked (Buxbaum et al., 2014); (3) background fluorescence is intrinsic to tissues and is aggravated by sub-optimal clearing of unbound probes from deep layers; (4) the refractive index can vary along the optical axis; (5) single-molecule signals must be identified in crowded environments and in varying image quality along the optical axis; (6) 3D-image reconstruction algorithms must work with anisotropic samples and (7) reliable segmentation of non-geometric, possibly incompletely labeled, 3D objects (e.g., nuclei) that extend over multiple 2D images in a discretely sampled image stack is required.

Here, we describe sample processing and imaging optimizations that have allowed us to perform smFISH in thick preparations of *Drosophila* larval body wall muscle. We optimized: (1) sample fixation and permeability conditions to improve probe penetration to deeper tissue sections and nuclei; (2) proteolytic processing to maximize mRNA accessibility to smFISH probes; and (3) time and temperature to improve probe signal and reduce background. We developed 3D-image analysis tools that allowed us to segment organelles and identify single-molecule signals in ~50 μm thick sections.

Using this optimized smFISH approach we investigated the composition and nuclear export of megaRNPs—giant ribonucleoprotein particles associated with the nuclear budding pathway for mRNA export. We find that nuclear mRNA complexes that could be interpreted as megaRNPs contain up to 62 copies of single mRNA species. MegaRNPs were only detectable inside nuclei, but not in the cytoplasm.

Although smFISH provides a versatile and accessible method for wide ranging inquiries into mRNA localization at the tissue level, data analysis becomes a limiting factor, especially if distances to organelle surfaces need to be measured with low bias. Oversimplified segmentation using symmetric shapes (e.g., boxes) for irregular shaped compartments (e.g., nuclei) is a major contributor to such bias. Our new 3D Gaussian fitting of smFISH data (3DISH) algorithm closes this gap.

## Results

To use smFISH to precisely localize individual mRNAs and megaRNPs within the cytoplasm and nuclei of *Drosophila* larval body wall muscles, we needed to optimize smFISH sample preparation for thick tissues, both to maximize signal and to minimize background throughout the tissue. Moreover, we needed to develop image analysis tools to automate spot detection, segment nuclei, and determine the positions of spots relative to one another and the nuclear membrane. We performed both in parallel, but for simplicity, we first describe our image analysis program—called 3DISH—followed by the smFISH optimization and our analysis of megaRNPs.

### 3DISH enables automated spot detection and organelle segmentation

Available spot analysis tools (e.g., FISH-quant and ImageM) provide different levels of 3D localization and precision, but no pre-existing tool could perform 3D segmentation by defining the perimeter or border of the nucleus as an irregularly shaped surface in 3D space (Lyubimova et al., 2013; Mueller et al., 2013; Raj et al., 2008; Semrau et al., 2013; Zenklusen et al., 2008; Zhang et al., 2013). We therefore developed a new 3D image-analysis tool that enables fast, parallelized 3D Gaussian fitting of smFISH data (3DISH).

3DISH supports automated 3D segmentation of nuclei using a continuous 4D hull function to fill gaps caused by discontinuous immunofluorescence staining of nuclear lamin, a marker of the nuclear periphery (**Fig. 2,3**). Physically collapsed or improperly segmented nuclei can be excluded from further analysis. Segmentation based on nuclear shape allows us to apply different parameters for spot detection in nuclear and cytoplasmic compartments and to independently account for different background levels and spot sizes. 3DISH also allows us to freely define 3D regions of interest (3D ROIs) and apply specialized image processing filters and spot detection criteria to each defined region (e.g., subtract local background, perform 3D spot detection, and verify spot location using 3D Gaussian model fitting and likelihood ratio testing). Gaussian fitting is slow, but graphics processing unit (GPU) implementation on a high-end gaming computer reduced the processing time to less than 10 minutes for an image stack of >50 planes and 1000x1000 pixels per plane. Finally, 3DISH allows us to calculate distances between individual spots or between spots and edges of segmented objects. We can therefore perform co-localization analyses using absolute 3D distances.

### Segmentation of multiple incompletely labeled organelles

Most image-analysis programs use box-shaped volume segmentation—i.e., a rectangular box drawn around the largest section of the nucleus and extending along the z-axis either throughout the whole image stack or to the top and bottom of the nucleus (e.g., **Fig. 2**). However, box-shaped volume segmentation misrepresents the curved shape of the nucleus and results in inappropriate projection of signals into the nucleus, even for signals in z-planes far above or below the nucleus. To distinguish between nuclear and cytoplasmic megaRNPs, we needed to segment nuclei according their exact envelope contours. We faced the following segmentation challenges: (1) Image stacks containing multiple nuclei; (2) individual nuclei extending over multiple z-frames; and (3) discontinuous nuclear lamin signal due to discrete z-sampling along the optical axis, nuclear invaginations, or incomplete immunofluorescence staining.

Segmentation (edge detection) by 3DG-FISH involves the following steps:

1. Two uniformly filtered image stacks—I_1_ (5×5×1-pixel kernel width) and I_2_ (10×10×1-pixel kernel width)—are computed from the original image stack (I_0_).
2. I_1_ is subtracted from I_2_, and the gradient of I_1_-I_2_ is computed, which results in a four-dimensional matrix of 3-dimensional vectors representing each pixel of the image stack (I_3_=I_1_-I_2_). The magnitude of each vector is calculated, resulting in again a 3-dimensional image stack (I_4_).
3. In I_4_, all small magnitude values, are limited to a minimum value max (I_4_,max(I_4_)/N), where N is user-defined. The parameter N depends on the signal to background ratio of the organelles that are to be segmented. In our case, N was typically 300 (arbitrary units).
4. A histogram-dependent, binary threshold (T_h_) is set on image I_4_ using the Isodata algorithm (Ridler and Calvard, 1978) resulting in I_5_= I_4_> T_h_. In the binary image I_5_, the value 1 represents volumes of the original image stack I_0_ for which a significantly high gradient was found.
5. To reduce false edges, the voxel numbers of separate objects are counted, resulting in a list with objects and their corresponding volumes. The mean volume of all objects in the image stack is calculated and all objects smaller than the mean volume size plus 1X the standard deviation of the mean are rejected.
6. The remaining edges can be discontinuous, caused by inhomogeneous staining of the nuclei. To connect the boundaries of individual nuclei, a dilation followed by an erosion is performed using a stencil with a width chosen by the user, based on the visual impression of nuclear sizes. The stencil size has to be chosen to be big enough to connect discontinuous edges but small enough not to connect adjacent nuclei. Often a stencil size of 10 is sufficient.
7. To identify the number of nuclei present in the image stack, the number of continuous volumes are identified and each volume is labeled with a unique identifier (Samet and Tamminen, 1988).
8. To segment the nuclei based on the size of the identified, possibly discontinuous edges, a three-dimensional convex hull is calculated by finding the smallest volume that includes all edges belonging to the same nucleus (prior assignment in step 6 (Barber et al., 1995)). Gaps are closed by linear connection of the end points, making it necessary that a sufficient number of possibly isolated edges are found during edge detection.
9. The resulting segmentation is used as an initialization point for 3D active contour optimization (Chan and Vese, 2001). In this optimization, the fitting energy of the contour is minimal when the curve is on the boundary of the object.
10. If the nuclear stain is found to be discontinuous, step 7 is repeated. For an example of segmented nuclei see **Figure 3**.

### Spot detection using local background subtraction

The major challenge for spot detection in thick tissues is accurate estimation of z-position. We needed to obtain an accurate z-position estimate to correctly assign RNAs as inside or outside of the nucleus and to measure the distance between RNAs and the nuclear envelope or RNAs and transcription sites. Inter-RNA distances are ultimately used to define colocalization criteria. In our experience, the most accurate methods for spot localization use a maximum likelihood-based (i.e., Gaussian) fitting approach (Deschout et al., 2014; Smith et al., 2010). For 3D images, a center of mass search can be combined with 2D-Gaussian fitting in the plane of the brightest voxel (Smith et al., 2015a). In thick samples, however, pinpointing the brightest voxel is limited by local background. Thick tissue samples are highly scattering, so the local background signal varies as a function of position along the optical axis. Inherent inhomogeneity of the biological sample—i.e., structures like nuclei or fibers—also contributes to local background. Fitting a 3D Gaussian is therefore more precise, as the z-position of a spot is not restricted by the discrete z-positions of image planes; 3D-Gaussian fitting uses the information in pixels across multiple image planes to find the best fit.

To identify smFISH signals in 3D, we applied an elliptic 3D maximum filter with parameters: 2*[σ_xy_ σ_xy_ σ_z_] to the background-corrected image stack (σ_i_ being the theoretical width of the expected PSF size). This filter is moved over all voxels in the 3D stack and creates a local maximum map. To find candidate positions for our Gaussian fitting routine, we searched the 3D local maximum map by comparing it to a 2D tophat background-corrected image stack. If a pixel had the same intensity value in the 3D maximum map and the background-filtered 2D tophat image, the pixel was accepted as candidate position. Dim candidate spots were rejected using an intensity threshold equal to the mean intensity plus N times the standard deviation of the background intensity in gray values. N was manually set to account for image quality—e.g., N=1 to 3 typically rejected non-specific smFISH probe signals. A different threshold can be set for each nucleus and cytoplasmic ROI, with the largest possible ROI being the whole image stack.

We centered each candidate position within a discrete candidate volume (an isolated cube) proportional in size to the theoretical expected PSF (3(2σ_psf_+1)) in pixels. For each cube, we extracted the deconvolved and aligned data and fit a 3D Gaussian, yielding the following parameters: x,y,z position, signal localization precision (σ_x loc_, σ_y loc_, σ_z loc_), intensity (I), background (bg), signal height (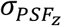) and width (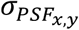), the false alarm probability (PFA), and the signal-to-background ratio: 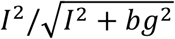.

These parameters can be used to manually define upper and lower limits, and control the number of false-positive detections. Because background is not uniform, parameter limits can be independently defined in the nucleus and in the cytoplasm for each channel. **Figure 3** shows a raw image (**Figure 3A**), the image after nuclei segmentation and mRNA detection (**Figure 3B**), and the images superimposed (**Figure 3C**).

### Single-molecule FISH optimization improves signal detection

To maximize the functionality of our 3DISH image analysis tools for smFISH in thick tissue samples, we also needed to optimize sample preparation and probe hybridization. Using published protocols as a starting point (Buxbaum et al., 2014; Lécuyer et al., 2008; Lyubimova et al., 2013), we systematically optimized sample preparation parameters (**Table 1**). We assessed the performance of each parameter change by examining the quality of anti-lamin immunofluorescence signal, nuclear morphology, and quantifying the ratio between double-labeled smFISH spots using two sets of probes, each labeled with a different color, and all spots (both double- and single-color) detected. Optimization for dual-color labeling of the same target, while maintaining immunofluorescence signal and nuclear morphology ultimately resulted in limited visual image quality while providing maximal information content.

**Table 1:** Optimization of sample preparation for combined smFISH and IF.

| *Table 1: Optimization of sample preparation for combined smFISH and IF* |  |  |  |  |  |
| --- | --- | --- | --- | --- | --- |
| **Fixation** | Concentration | Time | Temperature | Panel Figure 1 | Comment |
| Formaldehyde | 3.7% | 30 min | 20°C | B |  |
| Formaldehyde | 3.7% | 45 min | 20°C | A, C, D, E | **Best condition** |
| **Permeabilization** |  |  |  |  |  |
| Ethanol | 70% | 60 min | 4°C | C |  |
| Ethanol | 70% | 480 min | 4°C | Not shown | Number of smFISH signals reduced |
| Ethanol & Triton-X100 | 70% + 0.5% | 60 min | 4°C | Not shown | Sample damaged/fragile |
| Methanol | 70% | 2 min | -20°C | Not shown | Increased background fluorescence |
| Methanol | 70% | 10 min | -20°C | Not shown | Increased background fluorescence |
| Triton-X100 in PBS | 0.5% | 15 min | 20°C | A, B, D, E | **Best condition** |
| **Protease Digestion** |  |  |  |  |  |
| Pepsin | 0.01 µg/ml | 30 min | 4°C | Not shown |  |
| Pepsin | 0.01 µg/ml | 30 min | 20°C | Not shown |  |
| Pepsin | 0.1 µg/ml | 15 min | 20°C | Not shown |  |
| Proteinase K | 0.01 µg/ml | 30 min | 20°C | B, C, D, E | **Best condition** |
| Proteinase K | 0.02 µg/ml | 10 min | 20°C | Not shown |  |
| Proteinase K | 0.02 µg/ml | 30 min | 4°C | Not shown |  |
| Proteinase K | 0.03 µg/ml | 13 + 60 min | 20°C + 4°C | Not shown | Double incubation, sample damaged |
| **Post-Fixation** |  |  |  |  |  |
| Paraformaldehyde | 4% | 15 min | 20°C | B, C, D, E | **Best condition** |
| Paraformaldehyde | 4% | 30 min | 20°C | Not shown |  |
| **Post Fixation Permeabilization** |  |  |  |  |  |
| Triton-X100 in PBS | 0.1% | 15 min | 20°C | Not shown | No improvement |
| Triton-X100 in PBS | 0.2% | 15 min | 20°C | Not shown | No improvement |

**Table 2:** Final Protocol.

|  | Time [min] | Temperature | Step |
| --- | --- | --- | --- |
| 1 | 2 to 5 min per larvae | ~2°C (ice cold) | Dissect larvae |
| 2 | 45 | 20°C | Fixation with 3.7% Formaldehyde |
| 3 | 15 | 20°C | Permeabilization with 0.5% Triton X-100 |
| 4 | 30 | 20°C | Protease digestion with 0.01 µg/ml Proteinase K |
| 5 | ~3 | 20°C | Wash with glycine (2 mg/ml in PBS) |
| 6 | 15 | 20°C | Post-fixation with 4% paraformaldehyde |
| 7 | ~1 | 20°C | Wash with 0.1% Triton X-100 in PBS |
| 8 | ~1 | 20°C | Wash with 2xSSC containing 10% deionized formamide |
| 9 | 240 (4h) | 37°C | Hybridize smFISH probes and primary antibody, add 1 mg/ml tRNAs as competitor in hybridization buffer, final concentration |
| 10 | ~3 | 20°C | Wash with 2x SSC containing 10% deionized formamide |
| 11 | 60 | 37°C | Incubate with secondary antibody in the dark |
| 12 | ~3 | 20°C | Wash with 2x SSC containing 10% deionized formamide |
| 13 | ~1 | 20°C | Rinse with PBS |
| 14 | ~3 | 20°C | Mount with mounting medium |
| **Additional comments:** |  |  |  |
| • Penetration rate for fixative is ~ 1 mm/h (http://www.leicabiosystems.com/pathologyleaders/fixation-and-fixatives-2-factors-influencing-chemical-fixation-formaldehyde-and-glutaraldehyde/)<br>• Shorter initial fixation times lead to a reduction in number of detectable smFISH signals, longer post-fixation times (after permeabilization) also lead to a reduction in number of detectable smFISH signals<br>• Methanol permeabilization increase background levels to the point where no smFISH signals could be detected<br>• IF fluorescence staining did not compromise smFISH signal, e.g. due to RNase contamination of the antibody<br>• Hybridization specificity of smFISH probes improved in the presence of tRNAs but not with N20 DNA oligomers<br>• Increased hybridization temperature of 45 °C did not reduce non-specific smFISH signal, but did damage tissue<br>• Harsher washes with reduced 0.1x or 0.5x SSC, instead of normal wash with 2x SSC, did not reduce non-specific smFISH signal |  |  |  |

Duration of fixation affected the number and size distribution of faint smFISH spots, with a longer fixation time resulting in more homogeneous and robust signals within and between samples. Permeabilization was essential for probe and antibody penetration throughout the tissue, but excessive permeabilization reduced the number of smFISH signals, increased background fluorescence, or damaged the sample (**Table 1** and **Figure 1**). Protease treatment improved the accessibility of mRNA to smFISH probes in all but the harshest condition (Buxbaum et al., 2014). Compared to standard protocols, our optimized sample preparation for *Drosophila* muscle tissue significantly increased the number of mRNAs that could be successfully visualized by smFISH (**Figure 1**). Improvements were most pronounced in the nucleus, but were also observed in the cytoplasm. We conclude that the protease treatment and permeabilization not only resulted in better signals, but also in a better labeling of the mRNA population as a whole (compare **Figure 1 A,B** to **D,E**). Our findings emphasize that standard smFISH protocols may not be optimal for every tissue, so we recommend careful optimization of sample preparation conditions.

**Figure 1.**
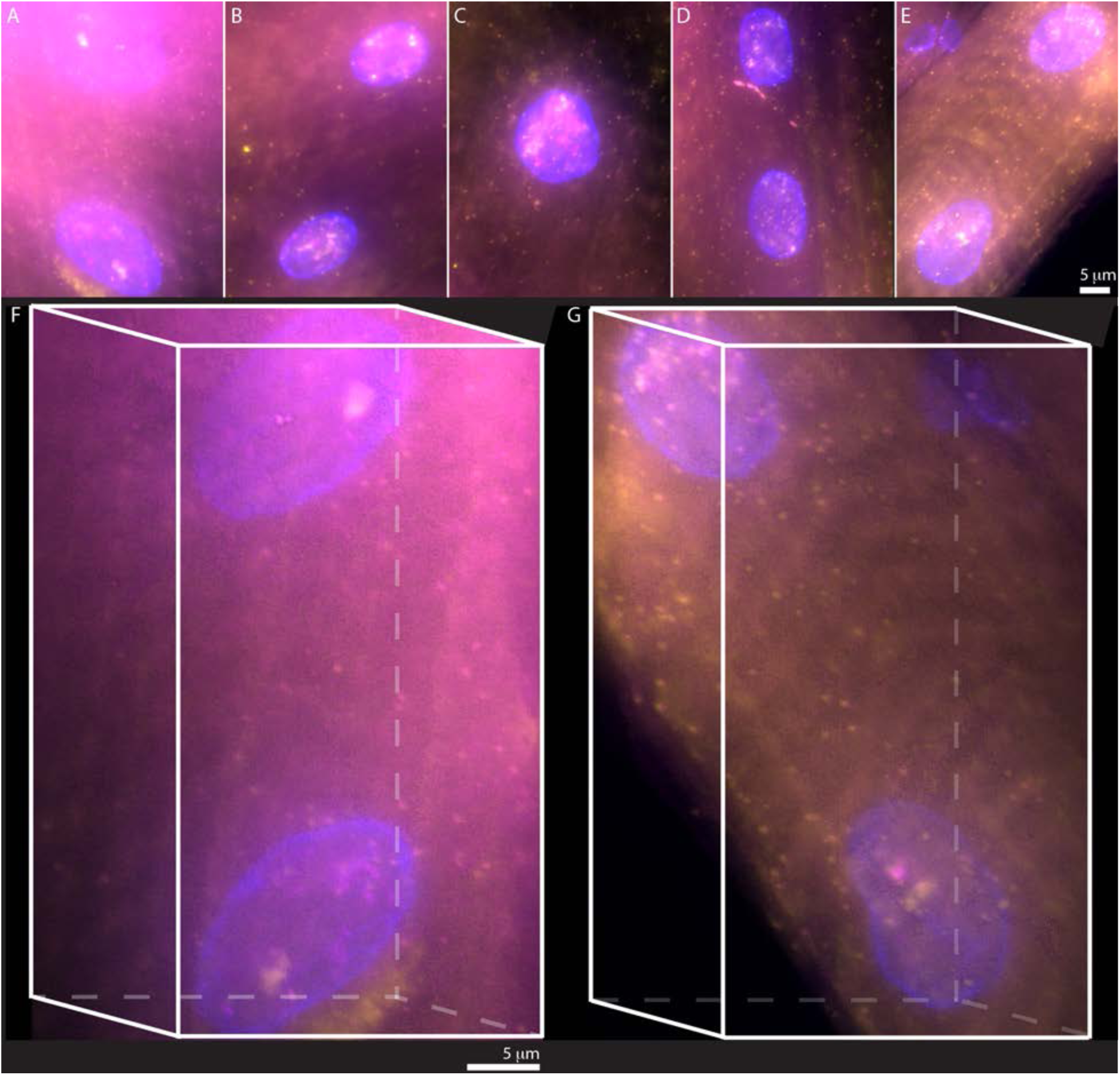
Protocol Optimization: The high amount of background and autofluorescence as well as limited accessibility of probes and antibodies to deeper tissue regions made it necessary to adjust existing smFISH protocols. A-E show maximum projections of 3D image stacks. A) Non-optimized protocol as provided by manufacture of FISH probes. B-E) Examples of optimized samples (see Table 1 for exact conditions for all panels). These are minor variations of the ideal protocol. To exemplify the variation between replicates: the sample shown in panel C differs from D&E (identical) only in the fixation step; samples in D&E are different replicates with strong differences in background. Based on our analysis of at least 3 biological replicates and at least 4 different fields of view per sample, we find that conditions D&E offer the highest reliability. F) 3D visualizations of A). G) 3D projection of E). Full stacks and 3D projections in supplement. Contrast and Brightness of all images were adjusted linearly to their minimum and maximum intensity, and the minimum value set above the background level. Green and red channels were assigned yellow and magenta lookup tables for visualization. Optimizations were done on >10 samples with >3 repeats for each protocol condition. Bar equals 5 μm, images were processed as described in the methods section.

**Figure 2.**
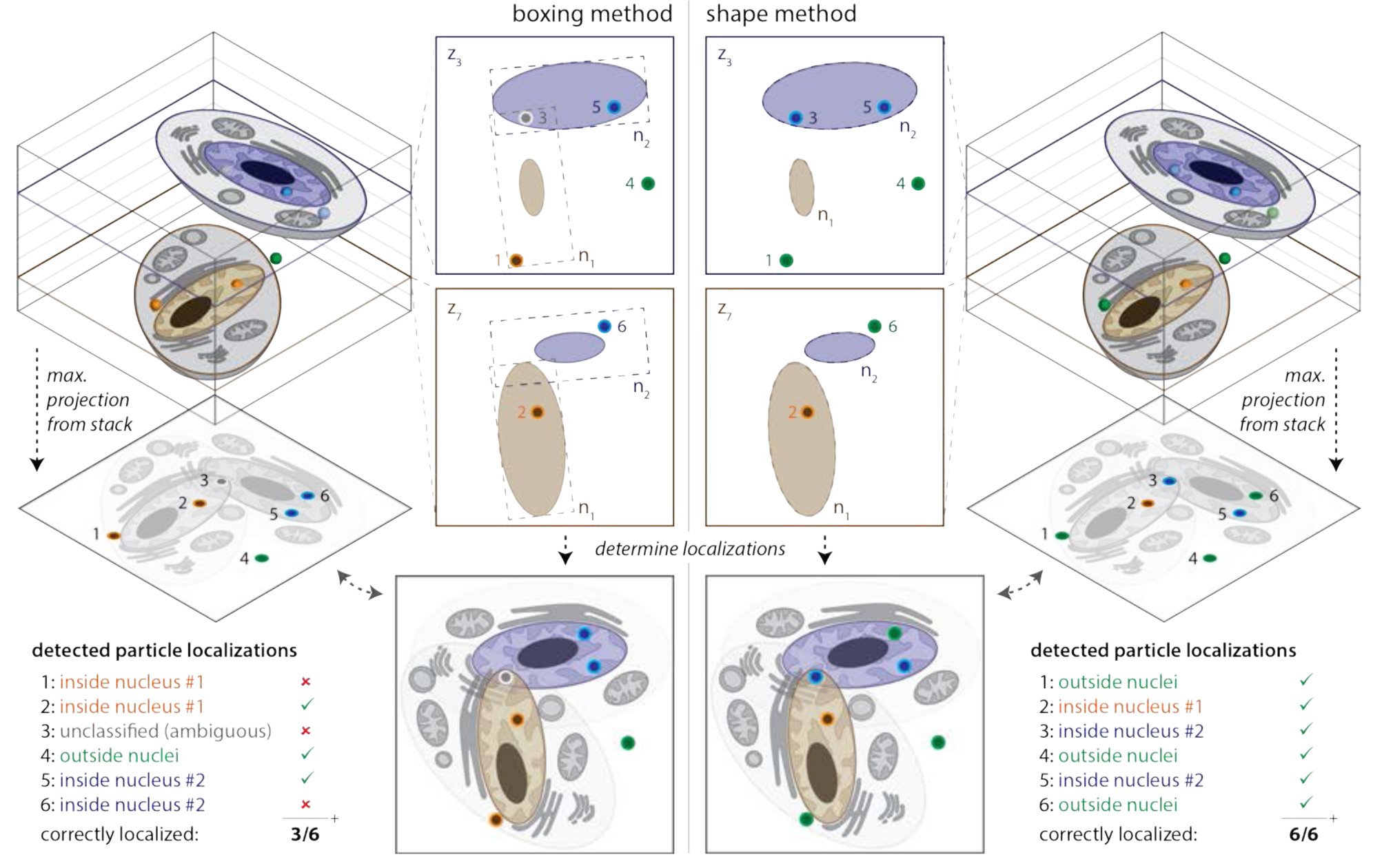
Relative localization of multiple signals in 3D: Left) Nuclei are segmented using a box (boxing method) that is set to enclose the structure in its mid-plane. This results in areas outside of the nucleus that are interpreted as being inside the nucleus (see panels for boxing method). If multiple nuclei exist in different z-planes of the image stack that overlap in the z-projection, boxing methods can lead to ambiguities when assigning smFISH signals to the correct nucleus. Right) If nuclei are segmented using a hull (shape method), the probability of correct assignment for smFISH signals in and close to the nucleus increases. The hull can also be used to estimate the shape of a nucleus that is incompletely labeled—often the case with lamin staining.

**Figure 3.**
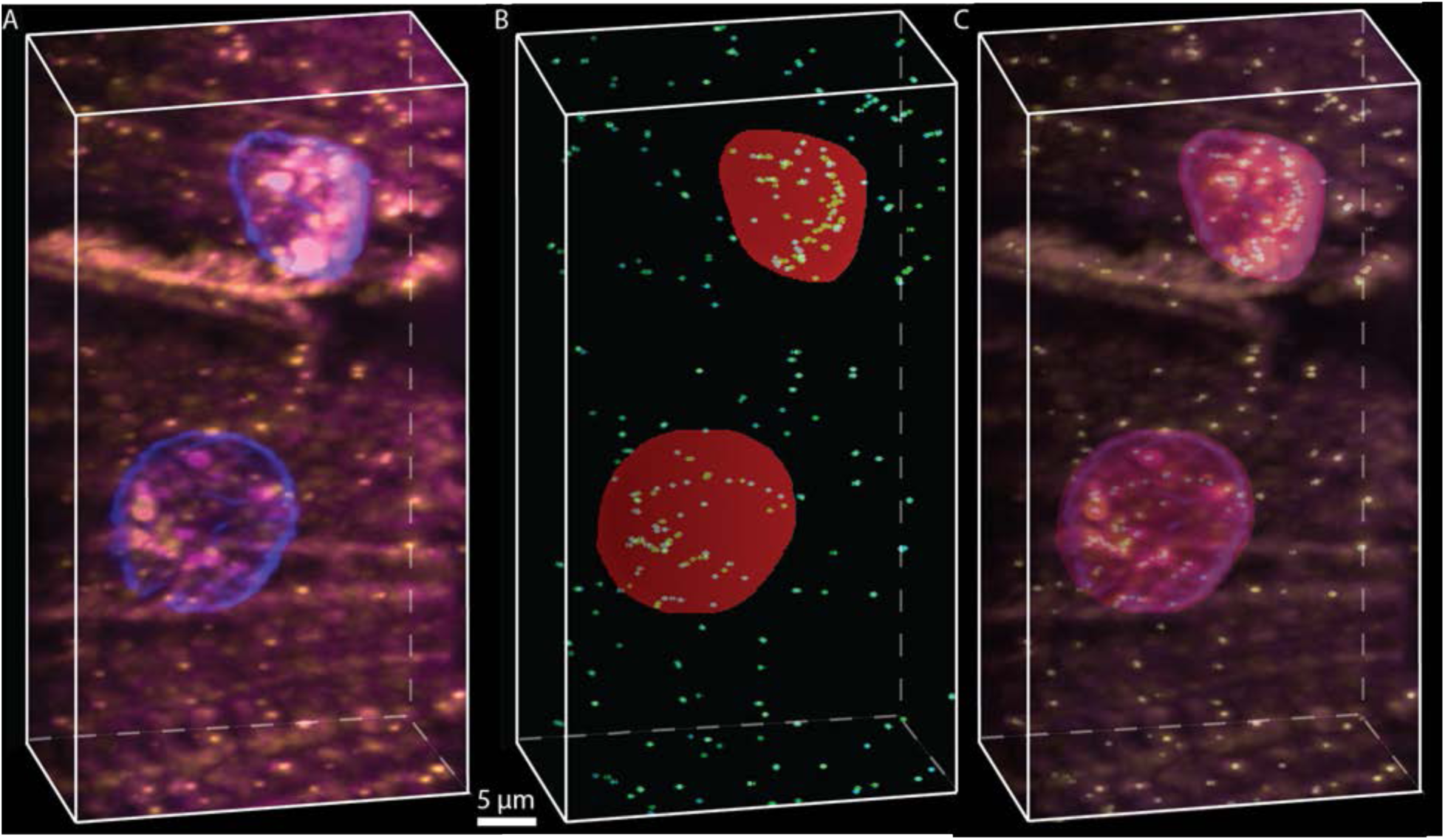
Automated nucleus segmentation and spot finding: A) Visualization of raw data for pak mRNAs labeled with odd (yellow) and even (magenta) probes. The nucleus is labeled with anti-lamin antibody (blue). B) Identical view of the 3D projected segmentation and spot finding results. Cyan indicates odd probes, green indicates even probes and red the nuclear volume. C) Overlay of A and B. Green and red channels were assigned yellow and magenta lookup tables for visualization.

**Figure 4.**
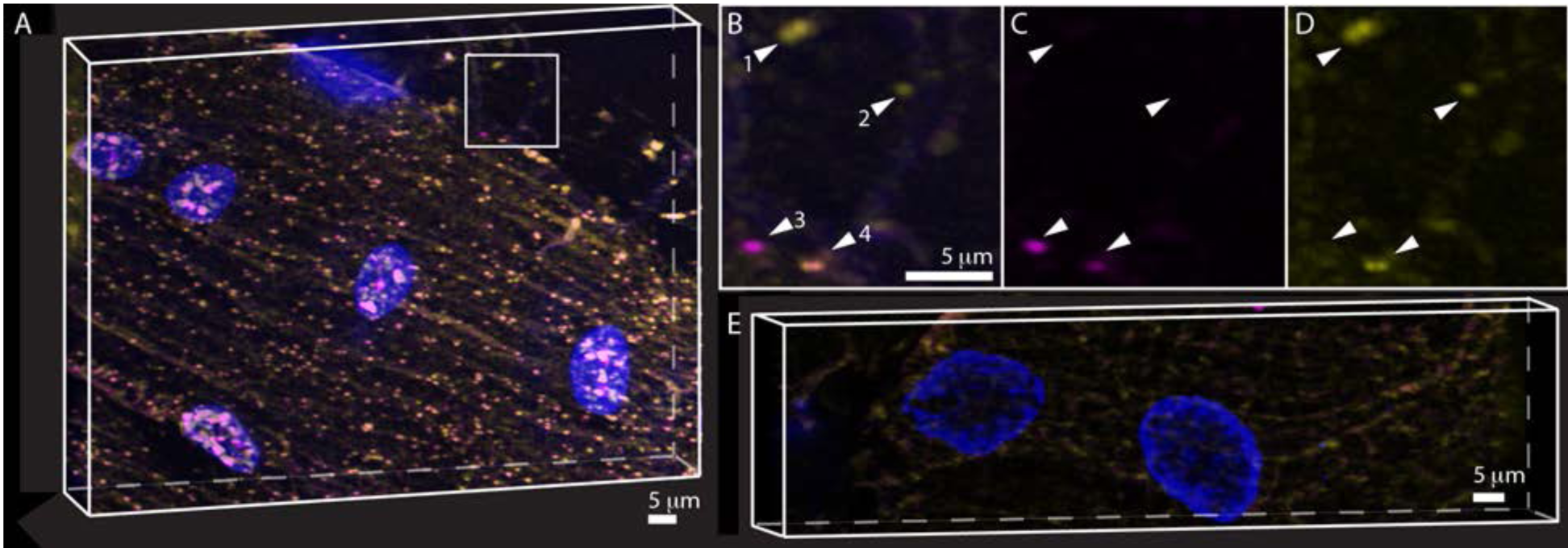
Probe specificity and reliability: Two probes sets with different dyes against Pak mRNA were used to test labeling efficiency through colocalization under ideal preparation conditions. A) 3D projection of one of seven replicates showing more than 90% colocalization. Arrows highlight typical examples of smFISH signals visible in both or only one channel. B-D) Zoom out from box in A), showing the colocalized (B), odd (C) and even (D) smFISH signals. Arrows as in A). Besides colocalization, the variance in signal between spots in each channel is noteworthy –see arrow 4 for example. Often a bright spot in one channel corresponded to a weak spot in the other channel. E) RNase treated sample processed (smFISH and IF) identically to A with equal intensity scaling. Contrast and Brightness of images A-D were adjusted linearly to their minimum and maximum intensity, and the minimum value set above the background level. Green and red channels were assigned yellow and magenta lookup tables for visualization.

**Figure 5.**
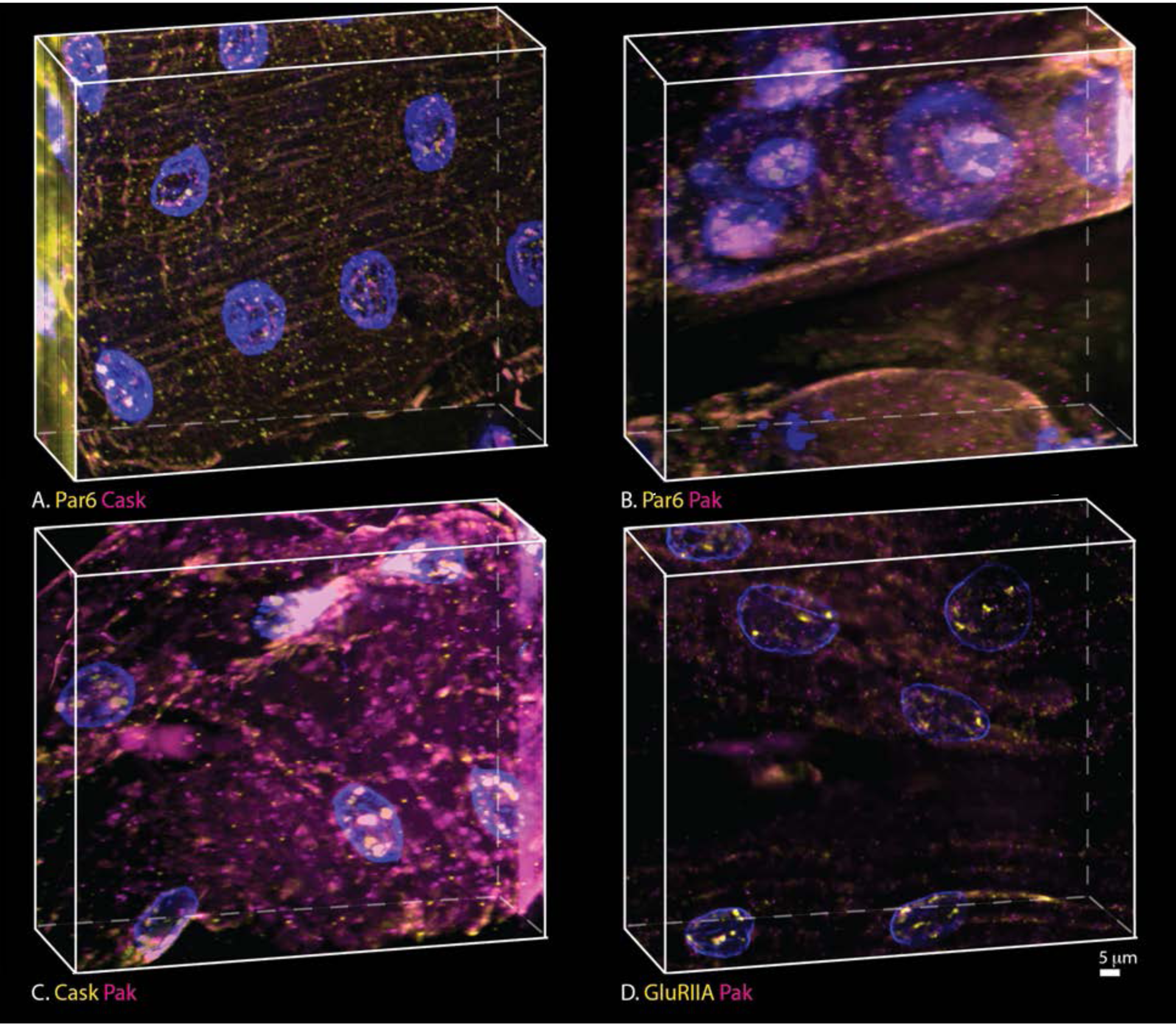
Composition of megaRNPs: Pak, Par6 and Cask mRNAs were detected in megaRNPs and predicted to travel together in megaRNPs. Colocalization analysis of smFISH for pairs of these mRNAs gives insight into whether megaRNPs contain multiple mRNA species. Nuclei are stained with anti-Lamin antibody (blue). A) Par6 and Cask mRNA smFISH; A) Par6 and Pak mRNA smFISH; C) Cask and Pak mRNA smFISH and D) Control using smFISH against GluRIIA and Pak two mRNA species not likely to colocalize. Green and red channels were assigned yellow and magenta lookup tables for visualization.

**Figure 6.**
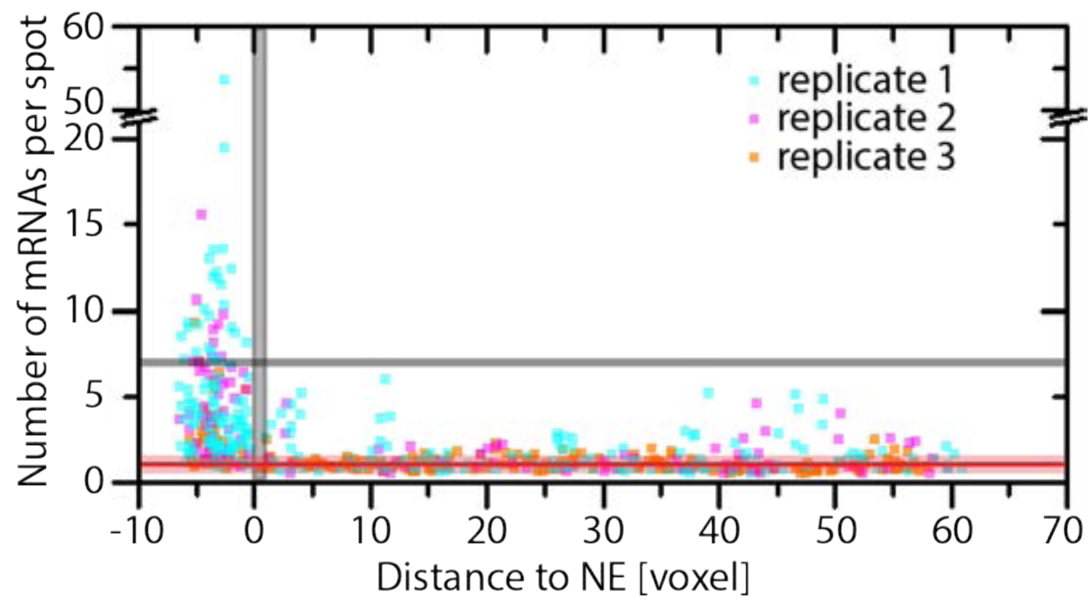
Distribution of megaRNPs: Brightness and distribution of double positive (odd and even probes) labeled Pak mRNAs. The vertical grey bar indicates the NE with a width of 2 voxels, note that in all data sets we find only one mRNA localizing inside the NE. the horizontal red line is set to 1 unit in normalized intensity with the transparent red bar indicating the 0.5 to 1.5 range of normalized intensity units classified as single mRNA. The horizontal grey line indicates an arbitrary lower threshold of 7 mRNAs per spot as cutoff between megaRNPs (>7 RNA) and single mRNAs or low copy number mRNA clusters. No megaRNPs are located in the cytoplasm. Cytoplasmic clusters of mRNAs with 5 or less mRNAs are more prevalent close to the NE. No data-points exist between 20 and 50 mRNAs. The data have been pooled from three replicates, which are indicated by color of the spots.

**Figure 7.**
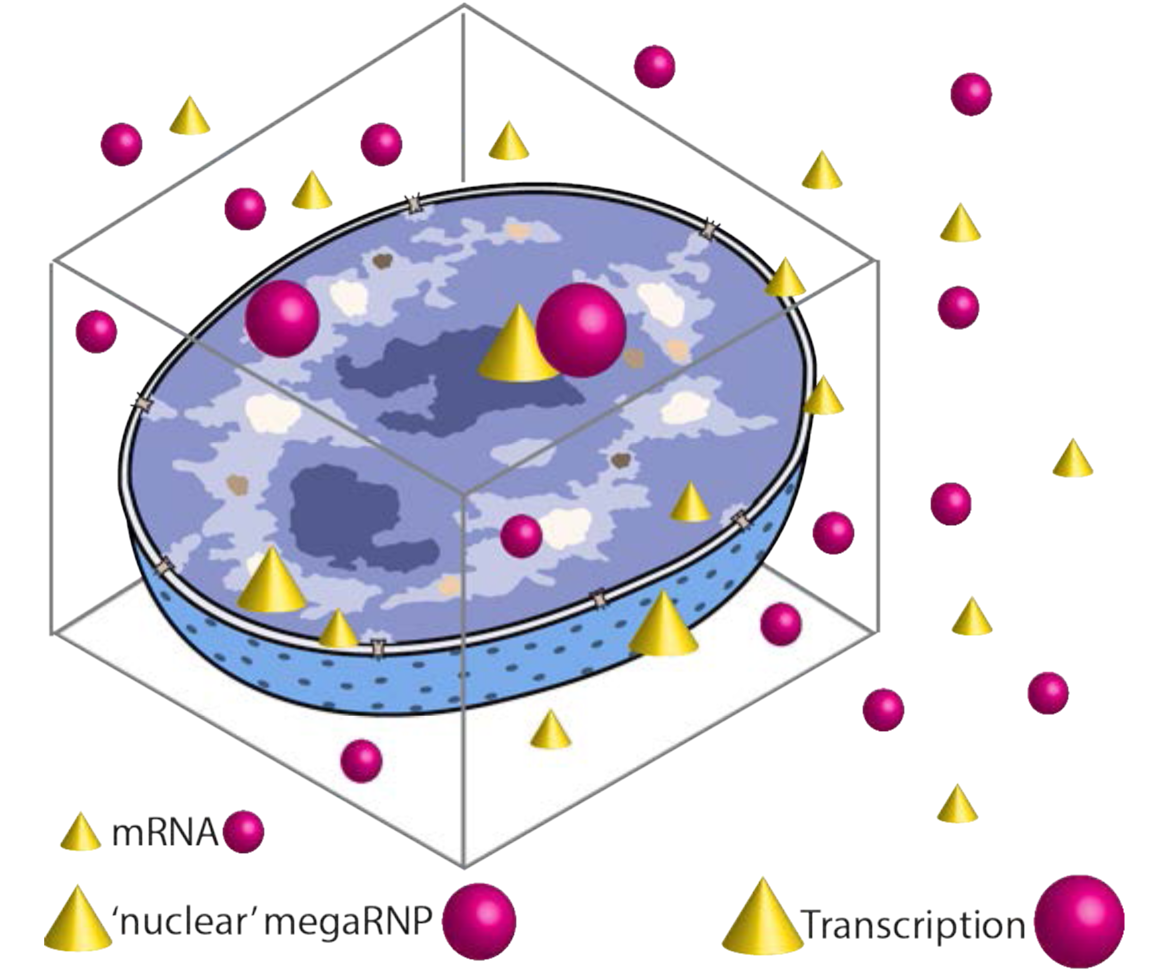
‘mega’RNPs at the single RNA level: For mRNA species previously identified as megaRNP components unusual large transcription sites exist in many nuclei. By the time megaRNPs locate to the nuclear envelope, single mRNAs of the same species can be found. Transitioning to the cytoplasm megaRNPs of Pak, Par6 and Cask likely distribute into single mRNAs.

Key to our optimization was our use of probes in two color spaces to label the *pak* mRNA. We designed 96 probes tiled along *pak* mRNA: 48 probes labeled in green (odd – visualized in yellow) alternating with 48 labeled in red (even – visualized in magenta). We did this in part because initial counting experiments labeling mRNAs with probes in a single-color space (e.g., 48 probes labeled in green tiled along *pak*, *par6*, or *cask*) yielded inconclusive or inconsistent results. We reasoned that probing a single mRNA species in two color spaces would increase confidence in the data. As a test, we imaged multicolor beads and achieved ≥90% colocalization using a spot-to-spot distance of ≤3 voxels (a voxel measures 107×107×200 nm) as the colocalization cutoff, as this is sufficiently large to account for chromatic aberrations. Prior to protocol optimization, only 36% of yellow and magenta *pak* smFISH spots co-localized in nuclei (N=3, range 25% to 49%) and 48% co-localized in cytoplasm (N=3, range 35% to 58%). Our optimized procedure increased co-localization frequencies in both compartments to 59% in nuclei (N=4, range 40% to 74%) and 72% in cytoplasm (N=4, range 63% to 81%). A strong smFISH signal in one channel was sometimes accompanied by a weak signal in the other channel (**Fig. 4 b-d**). Therefore, this weak signal would not be counted as a positive *pak* mRNA spot. smFISH signals detected by 3DISH were RNA-specific – ribonuclease treatment strongly reduced smFISH signals and virtually eliminated colocalization of yellow and magenta *pak* probes. Thus, despite extensive optimization of labeling conditions (see **Table 1** and **2**), probe accessibility to mRNA limits the smFISH labeling efficiency as can be seen by the low fraction (~80%) of dual color stained mRNAs. Nevertheless, using probes in two color spaces to label an mRNA does increase confidence in detected spots.

### Analysis of megaRNPs

MegaRNPs containing *pak*, *par6*, and *cask* mRNAs were previously observed at lamin C foci, in the lumen of the nuclear envelope, and at NMJs using digoxigenin-11-dUTP probes labeled with a-digoxigenin and secondary fluorescently labeled antibody (Speese et al., 2012). We sought to use smFISH to directly and simultaneously visualize *pak*, *par6*, and *cask* mRNAs in *Drosophila* muscle tissue. Because we never obtained 100% co-localization of yellow and red *pak* smFISH signals, however, spot detection of probes in one-color space could not be used to definitively accept or reject spots as an individual mRNA. But because each megaRNP likely contains many mRNA molecules (Speese et al., 2012), we expected that our assay would robustly report on the number and distribution of megaRNPs and whether an individual megaRNP contains single or multiple mRNA species.

MegaRNPs were proposed to export from the nucleus by budding through the nuclear envelope and to deliver mRNAs to the neuromuscular junction (NMJ) for localized translation. In tissues stained with *pak* probes, we analyzed the intensity and distribution of megaRNPs in nuclei and cytoplasm, accepting only double-labeled spots as true signal. Based on our data (**Fig. 6**) we defined candidate megaRNPs as spots with a minimum intensity seven times the average intensity of an isolated mRNA. Of 22 nuclei examined, 10 contained 2 or fewer smFISH signals meeting this minimum intensity requirement. Because transcription sites might also harbor multiple mRNA molecules, we excluded these nuclei from further analysis. The remaining 12 nuclei contained up to 37 candidate megaRNPs. Based on the brightness of isolated *pak* mRNAs in the cytoplasm, we estimated that the brightest nuclear megaRNP in our data set contained ~50 copies of *pak* mRNA (**Fig. 6**). This is likely an underestimate because labeling of mRNAs (especially nuclear) was likely incomplete, based on our 60% probability of detecting of *pak* mRNA, biasing the intensity analysis. Similarly, we often had to reject data that exceeded the dynamic range of our camera, especially for signals originating from nuclei.

Plotting normalized signal intensity (i.e., number of mRNAs per spot) against distance from the nuclear or cytoplasmic side of the nuclear periphery (**Fig. 6**), we observed a 2-voxel gap (~215 nm) in the distribution of spots, coinciding with the nuclear envelope (**Fig. 6**). Most cytoplasmic signals were between 0.5 to 1.5 normalized intensity, consistent with the detection of single mRNAs. Most of the brightest spots in the cytoplasm (1.6 to 6 normalized intensity) were within 2 μm of the nuclear periphery (**Fig. 6**). MegaRNPs were only identified inside the nucleus and were on average 11-fold more intense than isolated mRNA spots (range 7- to 54-fold brighter; median 9-fold brighter). The distribution of nuclear megaRNPs peaked at ~4 μm from the nuclear lamin signal, towards the center of the nucleus. Due to the limited number of megaRNPs, however, the distribution was not statistically significant. Therefore, in agreement with the original work (Speese et al., 2012), we found no strong correlation between the number of nuclear megaRNPs and distance to the nuclear envelope.

To address the question of whether individual megaRNPs contain one or multiple mRNA species, we performed smFISH experiments with differently colored probes for *pak*, *par6*, and *cask* mRNA, all of which are RNAs possibly forming megaRNPs, and *GluRII* mRNA (which was previously shown not to colocalize with megaRNPs; Speese et al., 2012). For our analysis, we take into account the known detection probability for a dual-color target from our optimization experiments using a dual-color probe set against *pak* mRNA. For *pak* mRNA we accepted only dual-color-labeled spots as true signal, while rejecting spots that were only visible in one of the two color channels, the detection probability, defined as the number of dual-color spots divided by all detectable spots, of *pak* mRNA was ~60% in nuclei and ~70% in cytoplasm. The presence of multiple mRNA species in a megaRNP will increase the probability of detection of megaRNPs above that of a single mRNA. For example, if a hypothetical megaRNP contained 3 copies of a single mRNA species the rate at which a megaRNP would not be detected is 6% and 0.001% for 7 copies – the miss rate for the megaRNP is given by the multiplication of the miss rates of the individual mRNAs. For our dual-color experiments for different RNA species we assume detection probabilities for each probe set similar to the *pak* mRNA probes and treat the observed binding of the single-color probes as an increase in the false-positive detection rate for megaRNPs. We did, however, not observe colocalization of *pak*, *par6*, or *cask* mRNAs in the cytoplasm any more frequently than we observed colocalization of *pak* mRNA with *GluRII* mRNA. Furthermore, colocalized signals of all possible pairs of those mRNAs were always of fluorescence intensity levels consistent with one or two mRNA molecules, not larger numbers (**Fig. 5**). Of 23 nuclei in which we detected both *pak* and *cask* signals, 9 nuclei had one observable *pak-cask* colocalization event and 2 nuclei had more than one. These isolated colocalization events could reflect proximity of the *pak* and *cask* transcription, which are located on the same chromosome.

## Discussion

In this study, we used single molecule FISH to investigate three mRNAs, *par6*, *pak*, and *cask*, that were previously identified as components of *Drosophila* larval body wall muscle megaRNPs (Speese et al., 2012). We aimed to better understand the composition and cellular location of megaRNPs. Single molecule FISH in thick tissue samples, however, presents numerous technical challenges. To overcome these, we optimized existing protocols and developed improved image analysis tools specific for 3D environments.

### Probe Specificity and colocalization calibration

Motivated by (1) initial probe colocalization rates far below our prediction (see **Fig. 1B**); (2) large numbers of presumably smFISH signals that yielded inconclusive results; and (3) the wish to employ probe mixes rather than synthesizing and optimizing each probe individually, we focused on finding a way to test and predict binding specificities of probe sets *in situ*. We worked with a mix of short, 20 nt probes as called for in newer smFISH protocols (Raj et al., 2008). These probes are cost-efficient compared to 50 nt or single sequence probes, allow for lower hybridization temperatures, and are often easier to use than longer probes that carry multiple dyes. In all, we designed a total of 96 probes against a single target mRNA (*Pak*). We numbered the probes sequentially in order of their location along the mRNA, and then divided them into two sets based on even and odd numbers. Odd and even probes were labeled with Quasar 570 and Quasar 670, respectively, yielding an interleaved pattern of red and green probes along the target mRNA. The probes had similar GC content and both sets were predicted to be of similar specificity to the target mRNA. We therefore expected that the colocalization frequency would only be limited by the optical performance of the microscope (>90% colocalization for multicolor beads), and not by the labeling protocol or probe performance.

A number of potential factors likely contributed to our initial lack of success using fixation, permeabilization, and protease digestion conditions previously developed for monolayer cells. For example, limited probe accessibility can lead to a scenario where the minimal number of probes needed for detection (estimated to be ~20; Raj et al., 2008) is reached after protease treatment for one probe set but not the other. In our case, however, colocalization rates did not increase upon changes to permeabilization conditions, protease digestion or hybridization times. If the number of colocalization events does not increase despite more protein being digested, probe accessibility is unlikely the reason the prediction and experiment fail to match. Single color, non-colocalized spots could be caused by properties of the dyes. To rule this out, we treated our tissue samples with RNase and found that all signals, both double-and single-color spots, disappeared (**Fig. 4**), indicating that they were RNA-dependent. Besides higher order folding structures of the target mRNA, this left open the possibility that substantial performance differences exist within our probe mix and that a subset of probes exhibited high-level off-target binding. We are unaware of any published data on the off-target probability of probes as a function of nucleotide length. It is worth noting, however, that smFISH was originally developed using 50-nt probes to increase probe specificity (Femino, 1998). A cocktail of 96 probes makes it difficult to test the efficiency of individual probes in the mix—either one by one or using established methods like ‘sense’ probes.

Despite these difficulties, we ultimately achieved sufficient probe performance to address several key questions regarding the formation and nature of megaRNPs. This is because the large number of mRNA molecules within megaRNPs minimizes detection sensitivity limits and averages out variations in probe accessibility to different mRNA regions. For example, if our colocalization-positive detection rate is 60% for a single mRNA, the overall colocalization detection rate increases to 99% if 5 mRNAs are present in a megaRNP (the miss rate in that case is 0.4^5^=0.01). While detection probability of megaRNPs is not limiting for our experiments, the variance in labeling efficiency introduces a bias for the intensity analysis of megaRNPs making the numbers of mRNAs per megaRNP a lower estimate. Probe specificity for colocalization experiments could be increased if each probe sequence was tested individually and a limited set of probes selected based on specificity and background performance. Such an effort might be useful if smaller mRNA complexes are targeted, but based on our ‘odd-even’ colocalization calibration we predict that the use of multi-color probes against the same target is a more viable strategy for assay optimization and to test overall assay performance.

### On the localization and composition of megaRNPs

Nuclear budding of large mRNP-granules, or megaRNPs was described as an alternative mRNA export pathway from the nucleus (Speese et al., 2012). Based on the mRNA binding factors found in such granules, it was hypothesized that a specific function of such megaRNPs is to promote the colocalized expression of functionally related proteins in specific locations in the cytoplasm (e.g., at a synapse). If this model is correct, then megaRNPs should be observable in both the nuclear and cytoplasmic compartments.

*<u>First,</u>* we investigated the hypothesis that individual megaRNPs contain multiple mRNA species by directly visualizing three different mRNA species (*Par6, Pak* and *Cask*) previously identified as megaRNP components (Speese et al., 2012). While limited to just these three mRNA species, our results favor a model in which megaRNPs are predominantly composed of a single mRNA species. This is consistent with data on neuronal mRNA granules where only single mRNA species were observed in dendritic processes (Batish et al., 2012; Knowles et al., 1996; Mikl et al., 2011), although reports of multiple mRNA species in mRNA granules do exist (Gao et al., 2008). The average number of mRNA molecules we find per megaRNP is ~10, but given our data and the physical size of megaRNPs (~200 nm; Speese et al., 2012) this estimate is, as discussed, a lower limit. *<u>Second</u>*, we investigated the composition of megaRNPs using smFISH, however, we were only able to observe megaRNPs within the nucleus. We therefore asked *<u>third</u>*, what the localization of megaRNPs in the differentiated cell would be if probed by smFISH. We based this analysis on smFISH using dual-color probes against *Pak* mRNA. The use of a dual color labeling of the same mRNA offers a higher degree of confidence in identifying ‘true’ mRNA signal if only dual-color spots are admitted for analysis. In our data, we find a clear two voxel wide gap between megaRNPs located in the nucleus and the cytoplasm as one would expect for nuclear pore complex mediate export in steady state cells. With an anti-lamin IF stain as label for the nuclear periphery we do not have direct knowledge of the position of the nuclear envelope, which in cells showing egress can be heavily displaced with the inner and outer membrane of the nuclear envelope being separated. While such separation should result in a wider range of distances without a clear local minimum we like to point out that budding of megaRNPs is likely a sporadic process possibly contributing to our results. We have only tested three examples of mRNAs we predicted to find in megaRNPs. The outcome for these three mRNAs differs clearly from our expectation but is limited to the mRNAs we tested. The use of single molecule FISH to study megaRNPs has several advantages as discussed in the introduction. We have limited our analysis to data sets with signal intensities within the dynamic range of our camera, but had to reject other data sets with higher signal intensity. We did so to be able to carefully control the technical quality of our work. For this very reason, we also optimized the labeling protocol using proteinase K and pepsin at various concentrations suitable for the study of RNA-protein complexes in neuronal cells. While the final protocol uses only 0.01 μg/ml protease, it is possible that the interactions between mRNAs and between proteins and mRNAs inside megaRNPs are very weak and lost in our experiments. While none of our controls or the visual impression of protease and pepsin free preparations – like Figure 1A, which is very similar with respect to the frequency with which we detect the bright megaRNPs – indicates so, we cannot rule out that single molecule FISH is less than an ideal technology to study megaRNPs. The image processing challenges in our work, however, are inherent to any 3D sample. The workflow and solutions we present here on the imaging aspects of detecting megaRNPs can easily be applied to other 3D experiments and introduce exact and defined measures for 3D distances and colocalization.

### Image Processing and Analysis

Image analysis of our 3D stacks, even after optimization of the smFISH protocol, provided challenges based on the thickness of the tissue for a number of reasons: (1) The refractive index of the sample changes with tissue depth. (2) As a result, signals from deep-tissue planes are weaker than from planes closer to the objective. The same effect was exacerbated by (3) the limited penetration of probes and reduced levels of protease digestion in deeper, more central tissue sections, (4) sample-specific background from filamentous structures within the tissue and (5) higher levels of background than found in single layer cultured cells. Another challenge was the need to minimize bias in locating smFISH signals inside or outside of the nucleus, and measuring the distance between a smFISH signal and the nuclear periphery. The origin of this challenge was, on the one hand the irregular shapes of nuclei - which leads to a high bias in ‘box’ approximations of their volume (**Fig. 2**) – and, on the other hand, the discontinuous labeling of the nuclear periphery by the lamin IF stain (**Fig. 3**). (7) The signal intensities we found for single mRNAs using a Delta Vision wide field microscope with deconvolution were too weak to be picked up reliably with the confocal microscopes available to us.

Lastly, robust detection of smFISH signals and estimating their 3D position turned out to be more complicated than we predicted based on the quantity of photons expected from the smFISH probes. This discrepancy is caused by variability in the number of probes bound per mRNA due to accessibility and specificity, as well as by optical aberrations and high background levels from the tissue. Here we described a method to segment nuclei by wrapping them in a hull that closely follows the shape of the lamin IF stain, to close gaps in this stain and in doing so largely eliminate any bias in compartment assignment. We found that this segmentation method also works well on other compartments; for instance, we have used it to identify neuronal junctions based on IF stains (data not shown). We combined this advanced segmentation with refractive index correction from optical disk drives (Smith et al., 2015a), true 3D Gaussian fitting of particle locations (Smith et al., 2010; Stallinga, 2005) and a maximum likelihood test to verify identified signals (Serge et al., 2008; Smith et al., 2015b). To reduce image processing time, we implemented graphic card processing (GPU) for 3D localization and likelihood testing. Despite all these advances, differences between compartments (e.g., different nuclei, nucleoplasm, cytoplasm, or other ROIs) were too extreme for fully automated analysis. We therefore enabled manual filtering options that could be individually applied to specific compartments. Combined with the ability to measure 3D distances between smFISH signals as well as distances between smFISH signals and any organelle surface, our analysis tool provides new and useful features currently not available to most biological research groups.

## Acknowledgements

We thank Adina Buxbaum and Robert H. Singer for sharing and helping with smFISH protocols, Vivian Budnik for discussions, reagents and critical comments on the manuscript, Vahbiz Jokhi for help with fly maintenance and Darryl Conte for editing and critical reading of the manuscript. We thank all RTI staff for their invaluable support. This work was funded by a Worcester Foundation grant and NIH grants R01 AI120902 and U01 EB021238 to D.G. and HHMI support to M.J.M.

## Author contributions

A.N., C.S.S., M.J.M and D.G. conceptualized the work; A.N., and C.S.S., performed formal analysis, validation, investigation and developed methodology with support from M.H., M.J.M. and D.G.; A.N. performed all biological experiments and protocol optimization. M.J.M. and D.G. administered and acquired funding for the project, C.S.S. developed software and algorithms with support from A.N.; A.N., C.S.S., M.H., R.M.M. and D.G. created figures; A.N., C.S.S. and D.G wrote the original draft; all authors reviewed and edited the manuscript. We used casrai.org contributor classification.

## Material and Methods

### Drosophila Stocks and antibodies

Wild-type flies (CantonS; CS) were raised at 25°C. IF staining was done using primary mouse anti-Lamin C antibody (LC28.26, 1:20-1:30, DSHB) and secondary anti-mouse antibody from Invitrogen (A31553, 1:200).

### Drosophila larva dissection

Third instar larvae were obtained and rinsed with ice-cold PBS. Larvae were dissected in ice-cold PBS, then immediately immersed in ice-cold fixation solution, see Table 1. Larvae were incubated at room temperature with gentle agitation, for times see Table 1. After fixation, larvae were rinsed three times with PBS, then kept in either 70%(vol/vol) ethanol at 4°C or PBS on ice until use. For RNaseA treatment, fixed larval filet was incubated with 100μg/ml RNaseA in 2XSSC at 37°C for 1hr then kept in either 70%(vol/vol) ethanol at 4°C or PBS on ice until use.

### Oligonucleotides

Single molecule Fluorescence In Situ Hybridization (smFISH) was done using Stellaris probes purchased from Biosearch Technologies (Novato, CA). All sequences are listed in Supplemental Table 1. Probe sequences were designed using the Biosearch Technologies online probe designer.

### Single molecule fluorescence in situ hybridization-immunofluorescence staining

Each probe pool against a specific mRNA contains up to 48 oligonucleotides with 20nt nucleotides in length (Supplemental Table 1). In situ hybridization was performed either by following manufacturer’s protocol or with modifications as specified in Table 1. Pepsin treatment was performed in 10mM HCl and proteinase K treatment was performed in 2XSSC. Pepsin and proteinase K were purchased from Sigma-Aldrich and Invitrogen respectively. After proteinase treatment, proteinase K solution was removed and quenched by 2mg/ml glycine in PBS. Larval filets were then washed 3 times with 2mg/ml glycine in PBS and post fixed as noted in Table 1. Larval filets were then washed twice with 0.1% PBT (0.1%(vol/vol) triton X-100 in PBS) and once with wash buffer for 5min or as shown in Table 1. Hybridization was performed for 4 hours at 37°C in hybridization buffer [10% (wt/vol) dextran sulfate; 2 x SSC 10% (vol/vol) deionized formamide; 1mg/ml yeast tRNA (Invitrogen), 2mM ribonucleoside vanadyl complex (NEB)] with 0.25 μM probe pool, with or without primary antibody. After hybridization, wash buffer was added to the hybridization solution. Then, larval filets were washed three times with wash buffer. For FISH-IF, incubation with secondary antibody was performed in hybridization buffer for 1hour at 37°C. Both hybridization and secondary antibody incubation were performed in microtubes with at least 50μl of reaction solution per 1 larval filet. Afterwards, larval filets were washed with wash buffer for three times, then three times with PBS and rinsed with water. Larval filets were mounted on a glass slide with Prolong Gold (Invitrogen).

### Image Acquisition

All images were taken with a DeltaVision Elite deconvolution microscope system (GE Healthcare Life Sciences), equipped with 60X/NA 1.40 oil immersion objective lens (PlanApo 60XO; Olympus) and a filter set to image fluorophores in DAPI, TRITC, and CY5 channels (center/bandwidth/nm: EX: 390/18, 542/27, 632/22; EM: 435/48, 594/45, 676/34), and a CoolSNAP HQ camera (Photometrics). The pixel size of the camera is 6.45 μm resulting in a pixel size in image space of close to 107 nm. The z-stack images were spaced 200 nm apart along the optical axis. Image size was cut to 1024 by 1024 pixels. Immersion oil (Olympus immersion oil AX9602) with a refractive index of 1.516 at room temperature was used. The embedding medium (Prolong Gold, Invitrogen) has a refractive index of 1.46.

### Deconvolution

A momentum-preserving deconvolution was applied and corrected for chromatic shifts (Parameter: area radius of background estimation 700 nm, CLME algorithm, stop criteria image differences <10^-5^ or 50 iterations maximum in Huygens, SVI Delft, The Netherlands). This deconvolution only relocates the photons and does not affect the actual photons counts. After deconvolution, the remaining local background signal was removed using a round (stencil) 2D tophat filter with a width of 8 pixels on each 2D image within the image stack, which was followed by segmentation of nuclei and spot detection.

In samples as thick as larval drosophila muscle tissue, variations in refractive index mismatch along the optical axis will contribute to a significant spherical optical aberration and wavelength-dependent defocus. The spherical aberration adds an additional source of image distortion and the z-shift creates a misalignment between the different channels. This situation is common when using high NA objectives, but due to the inherent 3D nature of our image acquisition, the spherical aberrations introduced on top of the expected defocus need to be taken into account in image processing (Smith et al., 2015a). The focal shift is caused by the wavelength dependency of the defocus function (Stallinga, 2005), which can be described as follows:

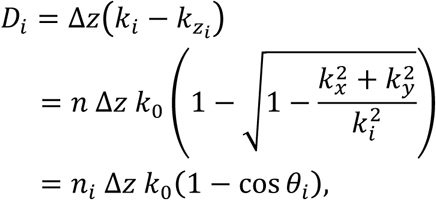

where n_i_ is the refractive index of medium i, k_0_ is the length of the wave vector in vacuum, k_zi_ is the wave vector length of the z component in medium i, Δz is the thickness of the medium. The differences in sign and constant phase offset to the original publication can be ignored. In addition to the original approach, we had to compensate for the spherical aberration introduced by refractive index mismatch. Furthermore, the spherical aberrations, also inherently dependent on wavelength, which is caused by index of refraction mismatch (Braat, 1997; Stallinga, 2005) and is given by:

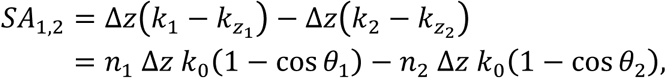

where Δz is the thickness of medium 2, θ_1_ is the angle in medium 1 and θ_2_ the angle in medium 2. Note that this expression does not only contain spherical aberration (of all orders) but also defocus.

First, the spherical aberration is estimated and deconvolution (Huygens, SVI The Netherlands) is performed with a z-dependent momentum preserving deconvolution to compensate for the depth dependent PSF distortion. Taking the spherical aberration into account for the deconvolution significantly reduces artifacts and increases image sharpness. Second, the different 3D color channels are aligned to adjust for the wavelength-dependent defocus, which we calibrated using tetra speck 0.2 μm beads (Invitrogen). Calibration was done by acquiring bead image stacks in all color-channels, deconvolving them and estimating the chromatic shift using the corresponding function in Huygens.

### Fitting of candidate spots

The likelihood of a signal measurement is assumed to be a Poisson process and is therefore given by:

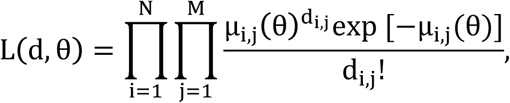

where μ_i,j_ (θ) is the expected photon count in pixel i, j and 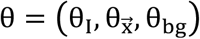 are the parameters to be estimated with I as the intensity, 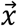 the three-dimensional position (*x*_0_, *x*_0_, *z*_0_).

The likelihood function is maximized by finding the zero of the negative log-likelihood function by use of Newton-Raphson. The derivative of the negative-log-likelihood is given by

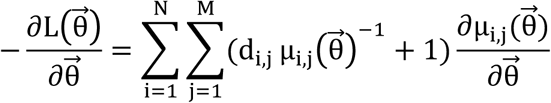

The precision with which we can determine the position of the single molecule depends significantly on the intensity of the single molecule, the background model, aberrations and the axial position; it is therefore essential to also estimate these parameters. This round of fitting is computationally expensive because a nonlinear optimization problem needs to be solved; the fitting was therefore done on a graphical processing unit similarly to (Smith et al., 2010), dramatically reducing the required computation time. The expected photons count depends on the choice of PSF. Here, the PSF is approximated by a Gaussian distribution:

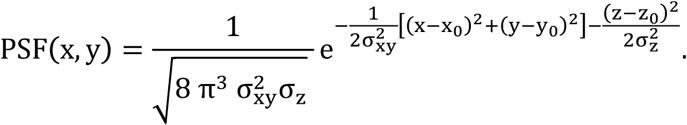

This PSF must be integrated over the pixel area to arrive at the expected photon count at each pixel k:

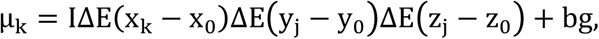

with:

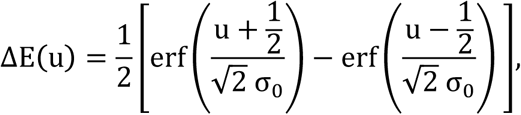

where (x_k_, y_k_, z_k_) are the pixel coordinates in unit (pixel) of pixel k, (x_0_, y_0_, z_0_) is the location of the center of the PSF in unit pixel and, σ_0_ is the PSF width, depending on the numerical aperture (NA), magnification (M), pixel size (Δp) and the wavelength of the light (λ).

The estimated parameters are total intensity (*I*), background intensity (*bg*), the PSF width (*σ*_*xy*_ and *σ_z_*), signal to noise ratio 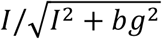 and the estimated Cramer Rao Lower Bound (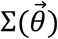) and the false alarm probability P_FA_ (Smith et al., 2015b).

### Distance calculation

Spot-to-spot distances were measured using the equivalent function of the MatLab tool box DIPImage (TU Delft, The Netherlands). Initially, the distance between a spot and a compartment edge where calculated by multiplying the compartment mask with a radial image (i.e. an image with the value of the R-coordinate of each pixel as the pixel values), and indexed the pixel in which an mRNA was detected. The most efficient implementation was accomplished by use of the k-nearest search algorithm presented in (Friedman et al., 1977), which finds the nearest periphery coordinate using a list of mRNA locations. This method provides us with the distance to the periphery with a maximum error of 1 diameter of a voxel.

